# Instantaneous amplitude and shape of postrhinal theta oscillations differentially encode running speed

**DOI:** 10.1101/2020.06.03.130609

**Authors:** Megha Ghosh, Benjamin E. Shanahan, Sharon C. Furtak, George A. Mashour, Rebecca D. Burwell, Omar J. Ahmed

## Abstract

Hippocampal theta oscillations have a temporally asymmetric waveform shape, but it is not known if this theta asymmetry extends to all other cortical regions involved in spatial navigation and memory. Here, using both established and improved cycle-by-cycle analysis methods, we show that theta waveforms in the postrhinal cortex are also temporally asymmetric. On average, the falling phase of postrhinal theta cycles lasts longer than the subsequent rising phase. There are, however, rapid changes in both the instantaneous amplitude and instantaneous temporal asymmetry of postrhinal theta cycles. These rapid changes in amplitude and asymmetry are very poorly correlated, indicative of a mechanistic disconnect between these theta cycle features. We show that the instantaneous amplitude and asymmetry of postrhinal theta cycles differentially encode running speed. Although theta amplitude continues to increase at the fastest running speeds, temporal asymmetry of the theta waveform shape plateaus after medium speeds. Our results suggest that the amplitude and waveform shape of individual postrhinal theta cycles may be governed by partially independent mechanisms and emphasize the importance of employing a single cycle approach to understanding the genesis and behavioral correlates of cortical theta rhythms.

## INTRODUCTION

Local Field Potentials (LFP) primarily represent summed transmembrane currents in the vicinity of an electrode and reflect neuronal ensemble activity (Ahmed and Cash, 2013; Buzsáki et al., 2012; Destexhe et al., 1999; Einevoll et al., 2013; Gold et al., 2006; Kajikawa and Schroeder, 2011; Katzner et al., 2009; Kelly et al., 2010; Telenczuk et al., 2017; Tingley et al., 2018). The information carried by LFPs is typically deciphered using frequency domain techniques that require averaging over several oscillatory cycles. Analyzing the frequency content of LFPs in this way has provided valuable information about neural oscillations and their aberrations in pathological conditions (Cunningham et al., 2006; Hutchison et al., 2004; Jensen et al., 2007; Lee et al., 2013; Lesting et al., 2013; Little and Brown, 2014; Marceglia et al., 2010; Neumann et al., 2014). Neural oscillations, however, change rapidly with time (Burns et al., 2011; Lopes-dos-Santos et al., 2018), with no two consecutive cycles being alike. Each single oscillatory cycle has instantaneous properties that can be quantified in the time domain, including amplitude, duration, nested oscillatory components, and waveform shape (Cole and Voytek, 2018; Gupta et al., 2012; Lopes-dos-Santos et al., 2018; Nelli et al., 2017; Zhang et al., 2019). These instantaneous cycle-by-cycle features can provide information that is complementary to that gained from spectral techniques (Amemiya and Redish, 2018; Cole and Voytek, 2018; Gupta et al., 2012; Lopes-dos-Santos et al., 2018; Nelli et al., 2017; Schaworonkow and Nikulin, 2019) and are the focus of the current study.

Hippocampal theta rhythms (~ 8 Hz) seen during active behaviors and REM sleep are among the largest amplitude oscillations in the rodent brain (Buzsáki, 2002). Hippocampal theta cycles transition into a temporally asymmetric, skewed, sawtooth-like waveform shape when an animal is running (Belluscio et al., 2012; Buzsáki et al., 1985; Sheremet et al., 2016; Terrazas et al., 2005). Although this asymmetry in the theta waveform shape is now known to be related to temporal computational dynamics such as phase precession (Belluscio et al., 2012), it is not completely understood. The relationship between instantaneous waveform shape (also referred to as instantaneous asymmetry) and other instantaneous features such as amplitude is not known. Whether this asymmetry in theta cycle waveform is seen outside of the hippocampus in other limbic cortical regions is also unknown. The postrhinal cortex (POR) is integral to the processing of spatial egocentric and allocentric information, as well as contextual information (Furtak et al., 2012; Aminoff et al., 2013; LaChance et al., 2019). Our previous work showed that strong theta rhythms are seen in postrhinal LFPs and a third of postrhinal neurons are phase-locked to theta (Furtak et al., 2012), indicating that theta rhythms influence local computations in this important brain region. However, the waveform shape of postrhinal theta rhythms has never been analyzed, so it remains unknown whether postrhinal theta cycles are asymmetric and whether their waveform shape changes with running speed.

Here, we develop improved instantaneous cycle-by-cycle approaches to understand the relationship between the amplitude and waveform shape (asymmetry) of individual theta cycles in the POR. We then use these methods to study how each of these instantaneous postrhinal theta cycle features (amplitude and asymmetry) correlate with running speed, which itself is a rapidly changing behavioral variable (Ahmed & Mehta, 2012). Using such instantaneous time domain metrics, we find that the falling phase of postrhinal theta cycles lasts longer than the subsequent rising phase, on average. However, the instantaneous amplitude and temporal asymmetry of individual theta cycles are poorly correlated. Furthermore, we show that the amplitude and temporal asymmetry of postrhinal theta cycles differentially encode running speed. Our results suggest that the amplitude and temporal shape of individual postrhinal theta cycles may be governed by partially independent mechanisms and emphasize the importance of employing a single cycle approach to understanding the genesis and meaning of rapidly changing cortical theta rhythms.

## MATERIALS AND METHODS

### Experimental Methods

#### Animals

Data collection methods are described in an earlier study that used this same data but did not analyze single cycle oscillation properties or asymmetry (Furtak et al., 2012). The current study is thus a secondary data analysis study. Briefly, for the first study, five male Long-Evans rats were used, and the same data was further analyzed here. Each rat was individually housed in a 12 hour light: 12 hour dark cycle. All behavioral experiments were performed between 10am and 4pm during the light phase. Before the start of behavioral training, animals were assigned a feeding schedule that helped maintain their body weight at ~90% of their free-feeding weight. All procedures were in accordance with the appropriate institutional animal care and used committee and NIH guidelines for the care and use of animals in research.

#### Surgery

Electrodes were implanted in the POR as delineated by Burwell (2001) and described in detail in our earlier study (Furtak et al., 2012). The implanted microdrive assembly was produced in-house and consisted of 8 individually movable stereotrodes (25 μm nichrome wires, A-M Systems, Inc., Carlsborg, WA). They were positioned at an angle of 22° along the mediolateral axis, 300 - 500 μm anterior to the transverse sinus, with their tips pointed in the lateral direction. Rats were trained after a recovery period of 7 days. Animals were given an overdose of Beuthanasia – D (100 mg/kg, i.p.) after the completion of the experiment. A small lesion was made at the end of each individual electrode tip position prior to perfusing the animals, extracting the brains, and performing histology. Subsequently, the locations of electrode tips were reconstructed with a light microscope and localized in the POR as defined by Burwell (2001). POR electrode locations are shown in our previous work (Furtak et al., 2012).

#### Task

The experimental setup consisted of an open field (81.3 × 81.3 cm) with images back-projected to the floor and the position of the animal tracked from above (Furtak et al., 2012). Food reward (chocolate milk) was delivered to four reward ports by computer-controlled pumps. Animals were trained on two discrimination problems (images), each consisting of a pair of high contrast, circular patterned stimuli. Animals were required to choose from this set of two different stimuli patterns. Food reward was delivered at the port behind the correct stimulus. Trials alternated between east and west. Each trial was divided into four epochs of 500 ms each: pre-stimulus (Ready), post-stimulus (Stimulus), pre-choice (Choice), and post-choice (Reward). A detailed description of the task can be found in Furtak et al. (2012).

#### Recording

Neuronal activity recorded from stereotrodes (McNaughton et al., 1983) was amplified (20x) at the head stage (HST/8o50-G20-GR, Plexon, Inc., Dallas, TX) and then passed through a differential preamplifier with a gain of 50 (PBX2/16sp-r-G50, Plexon, Inc.). LFPs were filtered between 0.7–170 Hz (PBX2/16sp-r-G50, Plexon, Inc.). The signals were then sampled at 1 kHz for LFP activity and further amplified for a total gain of 10,000 (MAP system, Plexon, Inc.).

### Data Analysis Methods

All time series and frequency domain analyses were performed in MATLAB. LFPs and timestamps were extracted using the FieldTrip toolbox (Oostenveld et al., 2011). One hundred and forty-five LFPs were analyzed for theta power ratio and extraction of individual theta cycles. The theta power ratio for a given LFP is the mean power from 6 to 10 Hz over the mean power from 4 to 12 Hz. The magnitude of net power can vary across electrodes due to factors such as the impedance of the electrode. Power ratios are often used to control for these differences in net power (Ahmed & Mehta, 2012; Cardin et al., 2009). Dividing theta power by power in the 4-12 Hz band and subtracting the baseline mean power from 0 to 40 Hz from both the numerator and the denominator allowed us to similarly normalize values across LFPs. LFPs were sorted by theta power ratio and divided into 3 groups based on increasing order of theta power ratio with cut-offs for each of the three groups determined by percentiles. Hence, LFPs in P1 had theta power ratios in the 0-33 percentiles, P2 had theta power ratios between 33-67 percentiles, and P3 greater than 67 percentile. Only P3 showed significant asymmetry in the theta waveform in this case (see Results). We obtained similar results if LFPs were divided into 9 groups based on percentiles with only the top three of the 9 groups showing significant theta asymmetry in this case (data not shown). Power spectral density was calculated using a wrapper function around Matlab’s built-in pwelch method of Fast-Fourier Transform (FFT) estimation (spectrum_pwelch).

#### Extraction of individual theta cycles

Local maxima and minima were found in the LFP filtered in the theta range. This helped to unambiguously identify a cycle that was centered on a trough. However, since the waveform shape is lost with the filtering, we additionally detected maxima and minima in a broader-band (6-40 Hz) signal in the cycle whose start and end times were defined by the previous step. Asymmetry analysis was performed on the individual 6-40 Hz filtered theta cycle extracted in this manner. This filter range (6-40 Hz) was empirically chosen to preserve the temporal shape of the theta cycle seen in the raw LFP.

### Asymmetry: Instantaneous Trough Location (iTL) Calculation

Trough location is the percentage of the total cycle duration spent on the downslope of the cycle. This was computed by dividing the start-to-trough time of a given cycle (trough is indicated by the purple dashed line in Figure 1) by the total duration of that cycle.

**Figure 1.**
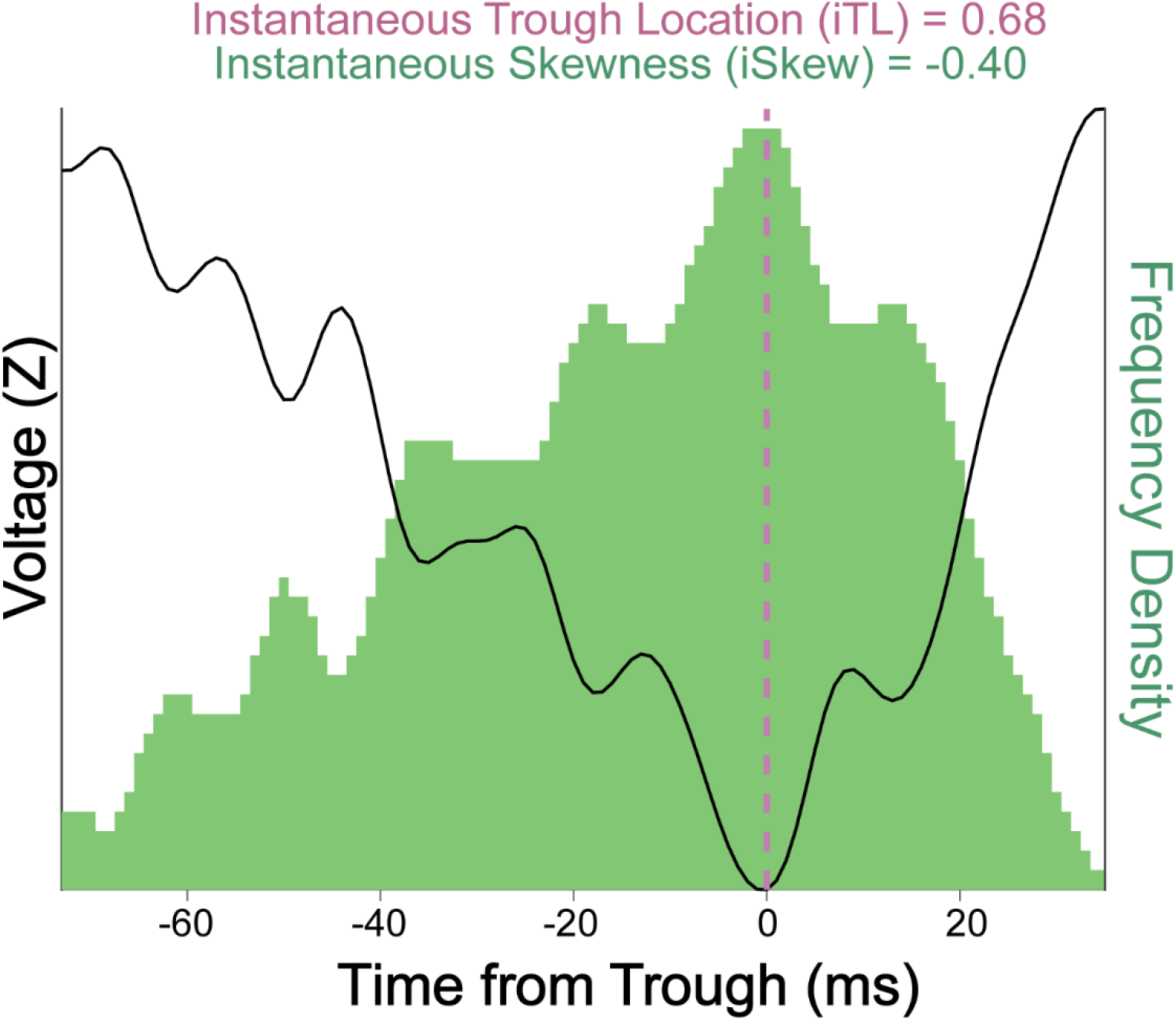
Two distinct methods to compute the asymmetry of a theta cycle. The black trace shows a single peak-to-peak theta cycle. The purple dashed line corresponds to the trough of the 6-40 Hz filtered theta cycle. We define instantaneous Trough Location (iTL) as the proportion of the total cycle duration spent on the downslope of the cycle. Thus, an iTL value of 0.5 corresponds to a symmetric cycle. The cycle shown here is asymmetric, with an iTL value of 0.68. The green histogram represents the cycle after it had been inverted, binned, zero-adjusted, converted into a frequency distribution, and displayed as a histogram (see Methods). We define the instantaneous Skewness (iSkew) of a cycle as the skewness of this inverted distribution (starting with a peak-to-peak theta cycle). By the definitions we use here, an iSkew value of 0 corresponds to a symmetric cycle. The inverted peak-to-peak cycle shown here is negatively skewed, with an iSkew value of −0.4. This corresponds to a theta cycle that has a slow voltage drop followed by a rapid rise. Note that if trough-to-trough theta cycles were used, then inversion would not be necessary, and the iSkew value would be positive.

### Asymmetry: Instantaneous Skewness (iSkew) Calculation

While trough location is an effective way to quantify the gross asymmetry between the duration of the falling versus rising phase of a theta cycle, it does not take into account differences in the precise shape and center of mass of each theta cycle. Asymmetry in any distribution can be quantified using the descriptive statistic skewness. For a single theta cycle, two types of skewness can be calculated: 1) skewness of the voltage distribution around the horizontal access (Bullock et al., 1997), and 2) temporal skewness around either the peak or trough of any given cycle. The latter definition of temporal skewness can capture the same asymmetry concept described by trough location and is the one we focused on in this study. We used instantaneous skewness (iSkew) as a novel measure of the temporal asymmetry of the theta cycle over time obtained by converting an individual cycle into a probability distribution. We used a peak-to-peak range to define one theta cycle because the peak-to-peak definition fits best with established analyses of theta phase precession (Buzsáki, 2002). Normally, using a peak-to-peak definition of a cycle would result in a bimodal distribution, so we inverted the values to obtain a unimodal distribution. To calculate iSkew, a theta cycle was first inverted and then digitized by multiplying it by an integer to space out the amplitude values (Figure 1). The cycle was zero-adjusted and incremented by one to prevent any bin from containing a value of zero. Each digitized value was stepped through with time as the x-axis. Each time value was appended to a new array ‘*y*’ times where y was the digitized LFP value (in arbitrary units; a.u.) at that time. For example, if *x* was 25 ms and *y* was 32 (a.u.), the value of 25 was appended to a new vector 32 times. This gave a vector containing the LFP represented as a frequency distribution. A histogram of this distribution was taken and used to determine the exact iSkew value (distribution in green, Figure 1). Since skewness is a well-established measure of asymmetry of any distribution, measuring the temporal asymmetry of theta cycles using iSkew allows the use of a standard statistical metric while capturing the true shape of the cycle at a finer resolution.

### Instantaneous Amplitude (iAmp)

The amplitude of each peak-to-peak defined theta cycle (iAmp) was computed as the difference between z-scored amplitude values of the starting peak and the trough, extracted from the 6-40 Hz filtered LFP.

### Calculating Speed and Testing for Significance

Animal position coordinates and the corresponding timestamps were loaded using a modified version of a Chronux function named readcont (original function is called nex_cont) that loads data stored in the NEX file format. We filtered the position (x,y) data and then used it to calculate speed of the animal obtained by 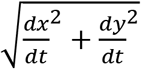. Since the position data was sampled at 100 Hz by the recording system, while the LFP data was sampled at 1 kHz, we interpolated the data using a shape-preserving piecewise cubic interpolation function (Matlab function interp1) to obtain the higher sampling rate. It was verified that the interpolation did not alter subsequent results.

### Statistics

A t-test was performed on the distribution of iTL and iSkew values of theta cycles to ascertain whether the cycles were significantly asymmetric (different from 0). Since an iTL value of 0.5 pertains to a symmetric cycle, 0.5 was subtracted from iTL values prior to the test. A Wilcoxon rank-sum test was used to compare squared asymmetry between each pair of three groups of LFPs, sorted by theta power (Bonferroni correction was applied). To understand the relationship of iAmp, iSkew, and iTL with running speed, theta cycles were divided into 4 speed groups (0-15 cm/s, 15-30 cm/s, 30-45 cm/s and >45 cm/s). A repeated measures ANOVA was performed on theta amplitude and asymmetry across these speed groups. Thereafter, post hoc comparisons using the Tukey HSD test were conducted. An alpha value of 0.05 was used across all tests.

## RESULTS

### Postrhinal theta cycles are temporally asymmetric

Analysis of the local field potentials (LFP) recorded from postrhinal cortical electrodes revealed that the theta waveforms were mostly non-sinusoidal and temporally asymmetric. Power spectral density (PSD) of the LFP recorded on a single electrode in the POR (Figure 2A) showed dominant power in the theta band (6-10 Hz) and high power in its second harmonic (12-20 Hz), which is often considered a signature of a temporally asymmetric waveform shape (Belluscio et al., 2012; Sheremet et al., 2016). This temporal asymmetry arises from theta cycles that are, when inverted and converted into a distribution over time, negatively temporally skewed on average. This means that the typical cycle takes longer to reach its trough than it does to return to the peak voltage (Figure 2B). We quantified this single cycle temporal asymmetry using two metrics: the instantaneous trough location (iTL) and a novel method that we call instantaneous skewness (iSkew; see Methods and Figure 1). A symmetric cycle, where the trough is midway between two peaks, is defined by an iTL value of 0.5 and an iSkew value of 0. The distribution of iTL (Figure 2C) and iSkew (Figure 2D) values for the example postrhinal LFP shown in Figure 2 was found to have a median iTL of 0.64 and median iSkew of −0.21, with both metrics indicating that these theta cycles were significantly skewed such that they had longer falling phases than rising phases (iSkew: t(11868) = −94.45, p < 0.0001; iTL: t(11868) = 91.3, p<0.0001)

**Figure 2.**
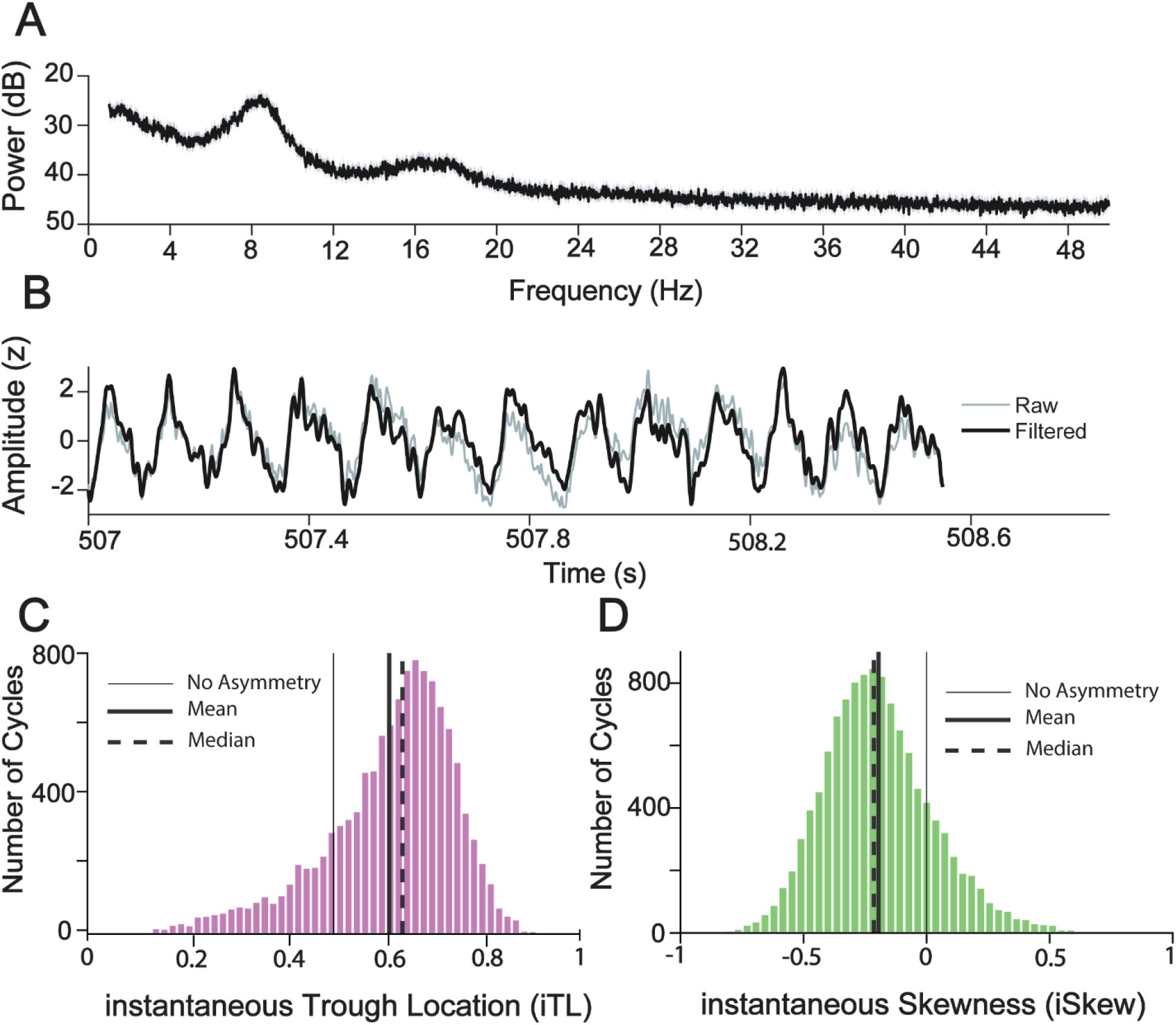
Postrhinal theta cycles are asymmetric. **A.** Power spectral density (PSD) of the local field potential (LFP) recorded on a single postrhinal cortical electrode. **B.** Example raw (grey) and 6-40 Hz filtered (black) postrhinal LFP recorded as the rat ran (speed >10cm/s). Setting the upper-end of the bandpass filter to 40 Hz preserves the shape of the waveform and reveals the clear asymmetry of these theta cycles. **C.** Instantaneous Trough Location (iTL), calculated for all peak-to-peak theta cycles in the same LFP. An iTL value of 0.5 represents a symmetric cycle, with the trough landing perfectly midway between the two peaks. The iTL distribution of all theta cycles (N = 11869, mean = 0.61, median = 0.64) in this postrhinal LFP was significantly asymmetric (t(11868) = 91.3, p<0.0001). **D.** Instantaneous Skewness (iSkew), calculated for the same peak-to-peak theta cycles analyzed in C. A value of 0 represents perfectly symmetric cycles, with center of mass at the midpoint of the cycle. The iSkew distribution of all theta cycles (N = 11869, mean = −0.20, median = −0.21) was significantly negatively skewed (t(11868) = −94.45, p < 0.0001).

### On average, the falling phase of postrhinal theta cycles lasts longer than the subsequent rising phase

Next, we examined the distribution of temporal asymmetry across all 145 postrhinal LFPs recorded across all electrodes and sessions. We sorted LFPs into 3 groups (P1, P2, P3; 48 LFPs in P1 and P3, 49 LFPs in P2) with increasing theta power ratio (Figure 3A,C; see Methods). To understand how temporal asymmetry changed across each group, we first characterized the magnitude of asymmetry by taking the squared values of the centered iTL and iSkew. The higher these values, the more asymmetric theta cycles are on average. We found that the magnitude of asymmetry of theta cycles was clearly highest in P3, the group of LFPs with highest theta power (Figure 3D, 3E). Statistical tests confirmed that the P3 group showed significantly higher squared asymmetry than either of the other two groups with lower theta power (Figure 3D, 3E; Wilcoxon rank-sum test, p < 0.001 in both cases, Bonferroni corrected). The fact that LFP locations where theta is essentially absent (e.g. P1) show virtually no asymmetry also indicates shows that asymmetry is not a random artifact of filtering: asymmetry only emerges when true theta cycles are present. Furthermore, as in the example shown in Figure 2, the asymmetry of LFPs selectively took the form of negatively skewed cycles at LFP locations with high theta power (iTL: t(47) = 5.8, iSkew: t(47) = −4.95, p < 0.001, Figure 3F, 3G). This is indicative of the falling phase of postrhinal theta cycles lasting longer than the subsequent rising phase. The LFPs in P1 and P2 were not significantly skewed in either direction (iTL: t(47) = 1.8, p = 0.06 and t(48) = 0.6, p = 0.52 for P1 and P2 respectively, iSkew: t(47) = 0.24, p = 0.8 and t(48) = 0.67, p = 0.5 for P1 and P2 respectively). This shows that as theta power increases, theta cycles in the POR become preferentially negatively skewed in the time domain. This is consistent with the fact that hippocampal theta oscillations also appear to be temporally skewed in the same way (Belluscio et al., 2012) and suggests that common mechanisms may regulate the waveform shape of theta oscillations across limbic brain regions.

**Figure 3.**
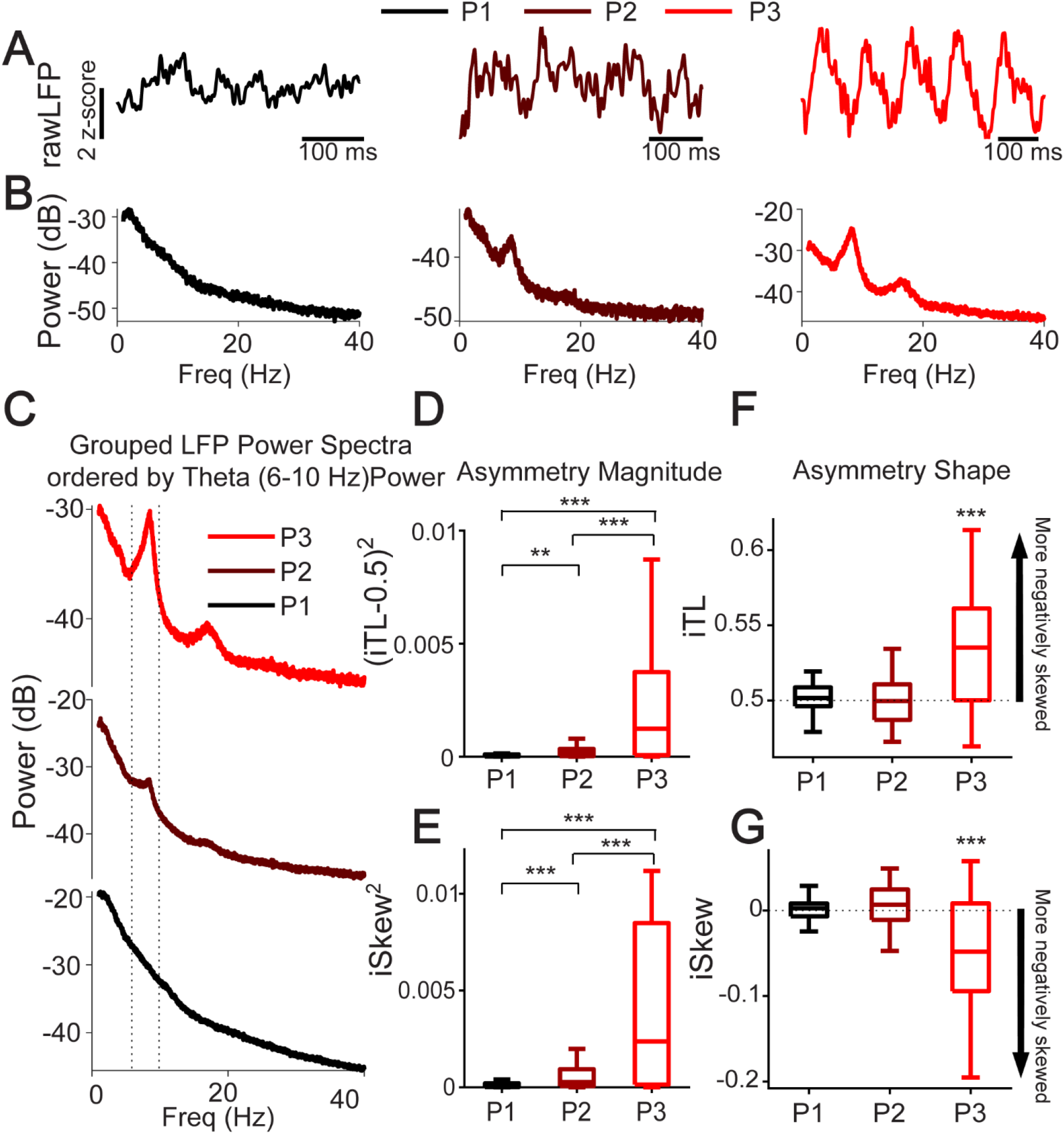
Postrhinal theta cycles are most asymmetric at locations with high theta power. **A.** One hundred and forty-five postrhinal LFPs recorded across all electrodes, sessions, and rats were sorted into 3 groups with increasing theta power ratio (P1, P2, P3). Representative LFP traces from each of the 3 groups. Running speed during each example was > 10 cm/s. **B.** Power spectra confirming the increasing theta power across the 3 representative examples from each group. **C.** The averaged power spectral density of the LFPs in each group, showing the increasing theta power from P1 (black) to P3 (red). **D.** The group with the highest theta power (P3) showed significantly higher asymmetry magnitude (represented here by squared centered iTL values) than either P1 or P2 (Wilcoxon rank-sum test, p < 0.001 in both cases; **: p < 0.01, ***: p < 0.001). Since a symmetric theta cycle has an iTL of 0.5, both positive and negative deviation from 0.5 would be indicative of asymmetric theta cycles. Squared centered iTL values allow us to establish the magnitude of this asymmetry. **E.** The group with the highest theta power (P3) showed significantly higher asymmetry magnitude (represented here by squared iSkew values) than either P1 or P2 (Wilcoxon rank-sum test, p < 0.001 in both cases). Since a symmetric theta cycle has an iSkew of 0, both positive and negative deviation from 0 would be indicative of asymmetric theta cycles. Squared iSkew values allow us to establish the magnitude of asymmetry. **F.** Panels F and G ascertain whether the asymmetry is negative or positive, giving more information about the nature of the waveform shape of theta cycles. Box-plot showing the distribution of instantaneous Trough Location (iTL) values in each of the 3 groups. iTL values were symmetric in groups with little to no theta power (P1, P2) but became significantly more asymmetric as average theta power increased (P3; t(47) = 5.8, p < 0.001). **G.** Box-plot showing the distribution of instantaneous Skewness (iSkew) values in each group. iSkew values were symmetric in groups with little to no theta power (P1 and P2) but became significantly more asymmetric as average theta power increased (P3; (47) = −4.95, p < 0.001)

### Instantaneous amplitude is uncorrelated to instantaneous temporal asymmetry of theta cycles

To understand the detailed relationship between theta amplitude and temporal asymmetry, we performed all remaining analyses on LFPs in the aforementioned group P3 with clear theta oscillations (48 LFPs; Fig. 3A). For any given LFP containing theta, if single cycle amplitude and asymmetry are determined by the same underlying mechanisms, then they should show a high correlation. To examine this correlation, we first plotted the instantaneous amplitude (iAmp; see Methods) against the instantaneous skewness (iSkew) of each theta cycle recorded on a given electrode. We found that iAmp and iSkew were essentially uncorrelated, with R^2^ values of less than 0.02 for each of the 3 examples shown in Figure 4, each for an LFP from a different rat (Figure 4A, 4B, 4C). The population data, plotted as the distribution of all R^2^ values across the 48 LFPs, showed the same result (Figure 4D). All of the R^2^ values were less than 0.12, and all but three were under 0.05. It is possible that the strongest correlations between amplitude and shape are only evident for the highest amplitude theta cycles in a given LFP. To test for this possibility, we repeated the correlational analyses, this time restricting the theta cycles to only the top 20% largest amplitude theta cycles on each electrode. Surprisingly, iAmp and iSkew remained weakly correlated even for these highest amplitude theta cycles (all R^2^ values were now less than 0.05; Figure 4E).

**Figure 4.**
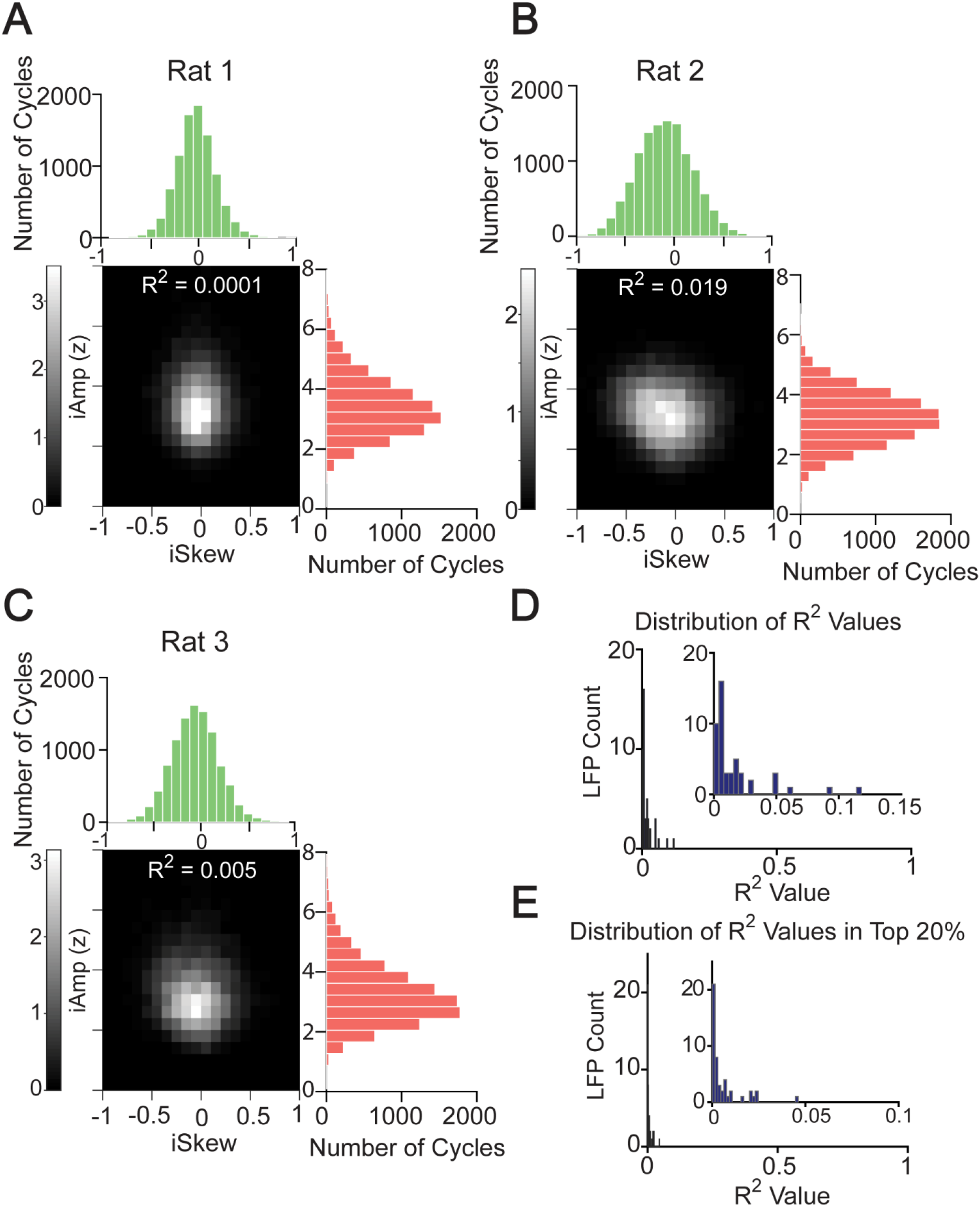
Theta cycle instantaneous amplitude and asymmetry are uncorrelated. **A-C.** Relationship between iSkew and iAmp for three postrhinal LFPs recorded in three different rats. The central density plot in each case shows the distribution of iSkew and iAmp for all cycles in a given LFP. iSkew and iAmp were uncorrelated in each of the examples shown here, with the R^2^ value being less than 0.02 in each case. **D.** R^2^ values for the relationship between iAmp and iSkew for all 48 LFPs analyzed. Inset zooms in to show that all R^2^ values fall below 0.12, and all but three values were below 0.05, indicating very poor correlation between theta instantaneous amplitude and shape. **E.** R^2^ values for the same analysis, now restricted to the 20% of theta cycles with the largest amplitude in each LFP. Once again, R^2^ values reflect very poor correlation between iAmp and iSkew with all values less than 0.05.

We next asked if the relationship between iAmp and iSkew might be stronger during specific behavioral epochs of the two-choice, visual discrimination studied in Furtak et al. (2012) or if the weak correlation was seen across all epochs of the behavioral task. Furtak et al. (2012) defined four behavioral epochs: ready (500 ms period immediately before stimulus presentation); stimulus (500 ms period immediately after stimulus presentation; selection (500 ms period immediately prior to a choice being made); reward (500 ms period immediately after the choice). We found that the R^2^ values were less than 0.12 in each of these task epochs (Figure 5A). These low correlations suggest that iAmp and iSkew are very poorly correlated within each behavioral epoch and hence potentially capable of carrying distinct behavioral information.

**Figure 5.**
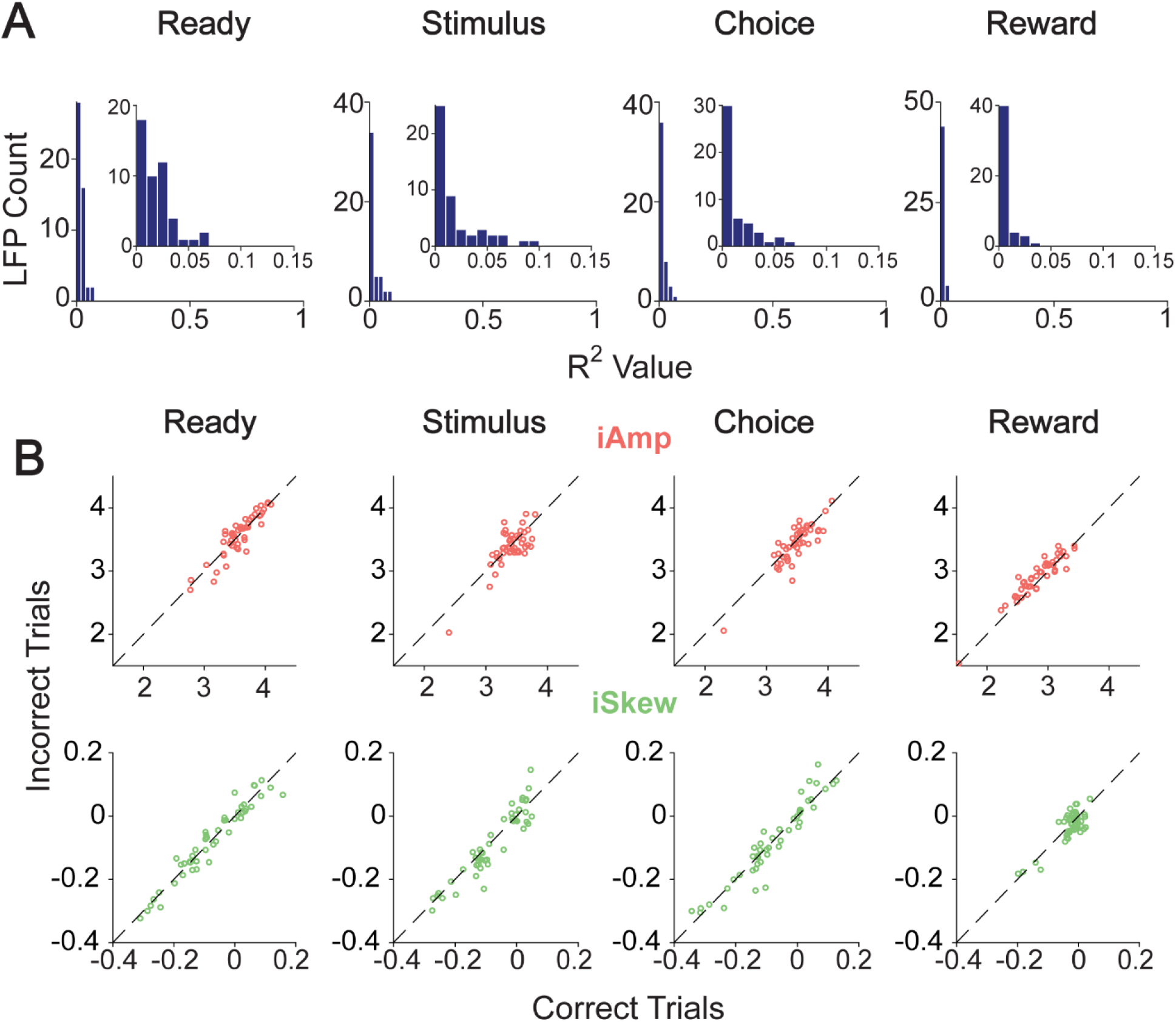
Theta cycle instantaneous amplitude and asymmetry are uncorrelated across all task epochs. **A.** R^2^ between iAmp and iSkew during the four task epochs (Ready, Stimulus, Choice, and Reward) across 48 LFPs. All R^2^ values in each task epoch fall below 0.12. **B.** iAmp and iSkew plotted for correct vs incorrect trials across all task periods; iAmp was significantly higher following incorrect (versus correct) choices during the reward epoch (Mean_correct_ = 2.8, Mean_incorrect_ = 2.9, t (47) = 4.7, p <0.001). All other comparisons were not significantly different.

Previous work on this dataset (Furtak et al., 2012) had reported that theta power, calculated using standard signal processing methods (FFT), was significantly greater following incorrect choices during the reward epoch. So, we asked if measures of instantaneous amplitude (iAmp) and instantaneous shape (iSkew) were also different during incorrect vs correct decisions across the task phases (Figure 5B). We found that similar to theta power, iAmp was higher following incorrect choices during the reward epoch (Mean_correct_ = 2.8, Mean_incorrect_ = 2.9, t (47) =4.7, p <0.001). Importantly, iSkew was not significantly different after correct vs incorrect choices during the reward epoch, again highlighting the ability of amplitude and shape of theta cycles to encode different information. This difference furthers emphasizes the independence of theta amplitude and asymmetry.

### Instantaneous amplitude and shape of theta cycles show distinct relationships with running speed

Postrhinal theta power (as measured using FFT) increases with running speed (Furtak et al., 2012). We next used single cycle methods to ask whether the same relationship is obtained when using the instantaneous amplitude (iAmp) of each theta cycle (Figure 6). Indeed, we found that theta iAmp increased as a function of increasing running speed in all rats and in the population averages (48 LFPs, Figure 6B, 6C). We next repeated the same analysis, but for the instantaneous shape of postrhinal theta cycles. We found that while the instantaneous cycle amplitude increased almost linearly with increasing running speed, both temporal asymmetry metrics (iSkew and iTL) peaked at medium speeds (around 25-30 cm/s) and then remained roughly the same at faster speeds (Figure 6D, 6E). The population averages showed similar saturating relationships for both iSkew and ITL.

**Figure 6.**
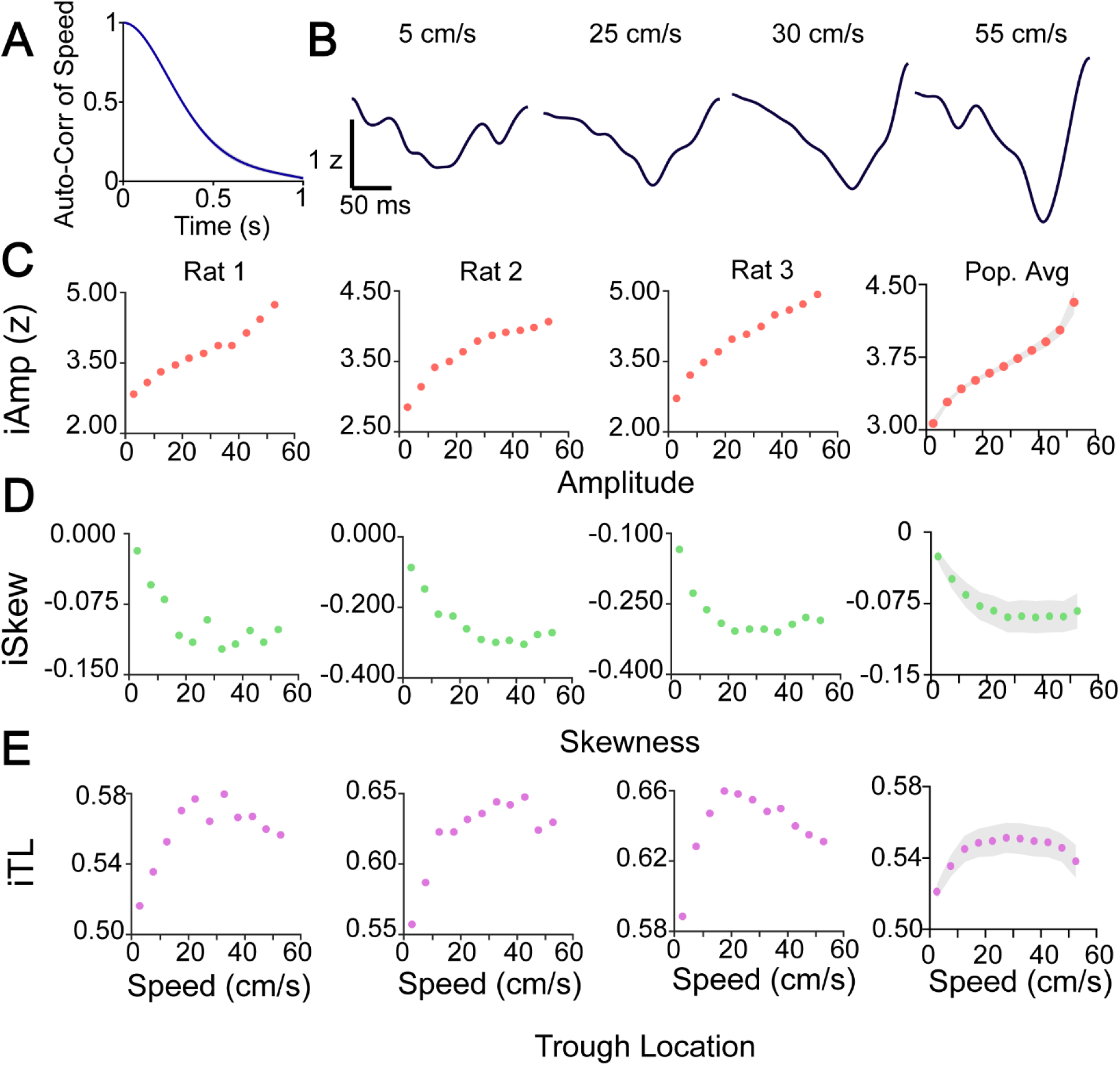
Postrhinal theta cycle amplitude and asymmetry show differential relationships with running speed. **A.** Auto-correlation of speed across the population (21 sessions). **B.** Representative theta cycles extracted from an example LFP but occurring at difference running speeds. Cycles had been filtered in the 6-40 Hz range. **C.** Variation of iAmp with running speed across three rats along with their mean values across the population. 48 LFPs were used to compute the averages across rats. The standard error for each speed bin is shown using a light grey shading. **D and E.** Change in iSkew and iTL with running speeds in 3 example rats and across the population. While amplitude increased with increasing speed, both asymmetry indices initially increased with speed and then quickly saturated around 25-30 cm/s.

To better assess this relationship between speed and single cycle features, we grouped cycles into four equal-sized speed groups spanning the full range of speeds sampled by the rats (0-15 cm/s, 15-30 cm/s, 30-45 cm/s and >45 cm/s). A repeated measures ANOVA was conducted to compare the effect of speed on instantaneous cycle properties (Figure 7). There was a significant effect of speed on iAmp (F(3,141) = 76.7, p<0.0001). Post hoc comparisons using the Tukey HSD test indicated that the mean iAmp for the 15-30 cm/s group (M = 3.58, SD = 0.17) was significantly higher than the 0-15 cm/s group (M = 3.15, SD =.28). Similarly, the mean iAmp for the 30-45 cm/s group (M = 3.8, SD = 0.27) was significantly higher than the 15-30 cm/s group, and the mean iAmp for the >45cm/s group (M = 4.1, SD = 0.48) was significantly higher than the 30-45 cm/s group. This confirmed that iAmp increases consistently across all speed bins.

**Figure 7.**
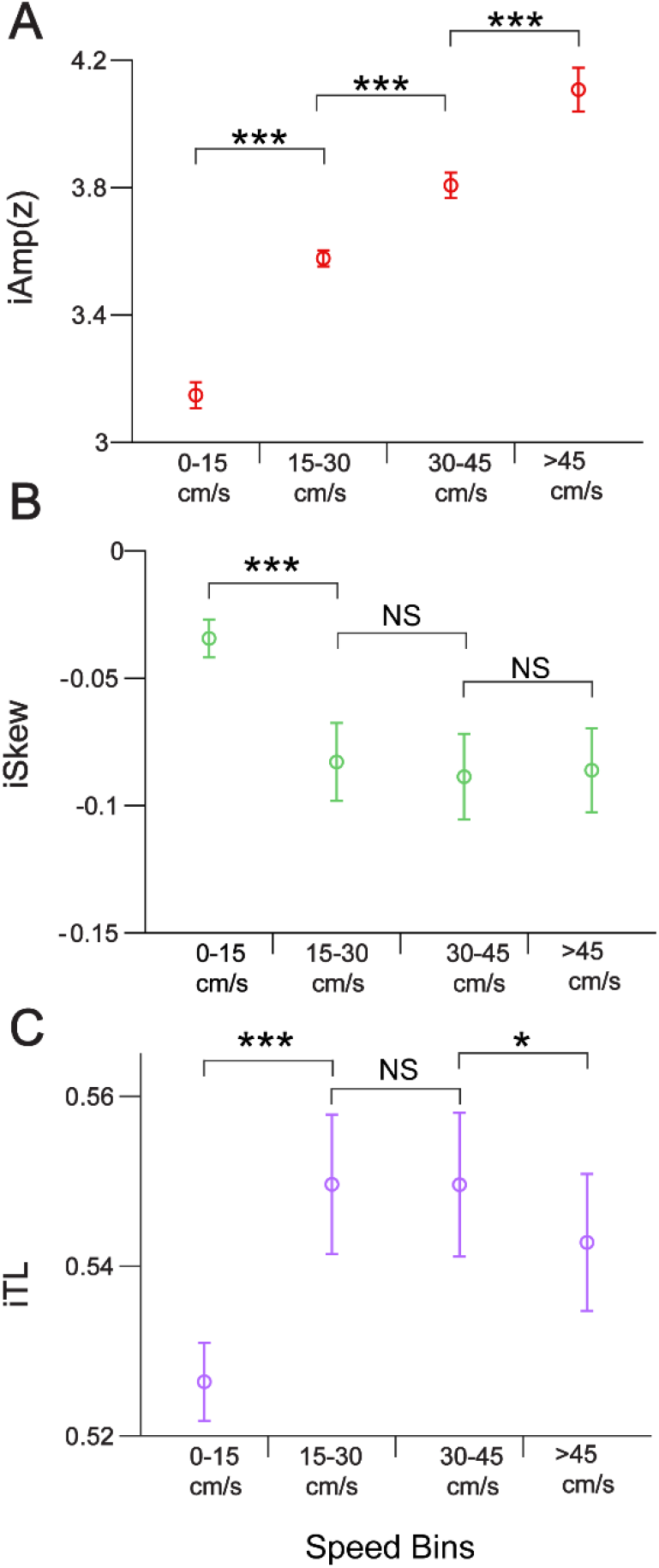
Instantaneous theta cycle amplitude, but not asymmetry, continues to increase at faster running speeds. **A.** Mean iAmp and standard error of theta cycles (48 LFPs) with increasing speed. The entire speed range was divided into 4 groups: 0-15 cm/s, 15-30 cm/s, 30-45 cm/s, and >45 cm/s. A repeated-measures ANOVA showed that there was a significant effect of speed on iAmp (F (3,141) = 76.7, p<0.0001). Post hoc comparisons using the Tukey HSD test indicated that iAmp was significantly higher for a speed group compared to every group with slower speeds. This confirmed that iAmp continues to increase with running speed, even at fast speeds (***: p < 0.001, **: p < 0.01,*: p < 0.05). **B.** Similar to **A**, but for the temporal skewness measure, iSkew. There was a significant effect of speed on iSkew (F(3,141) = 24, p<0.0001). Post hoc comparisons using the Tukey HSD test indicated that the mean iSkew for the 15-30 cm/s group was significantly more negative than for the 0-15 cm/s group, indicative of increasing temporal asymmetry at 15-30 cm/s compared to slower speeds. However, the mean iSkew for the 30-45 cm/s group was not significantly different from either the 15-30 cm/s or the > 45 cm/s groups, indicating that the temporal asymmetry of theta cycles quickly reach a plateau at medium speeds. **C.** Similar to **A**, but for the temporal skewness measure, iTL. There was a significant effect of speed on iTL (F(3,141) = 19.8, p<0.0001). Post hoc comparisons using the Tukey HSD test indicated that the mean iTL for the 15-30 cm/s group was significantly higher than for the 0-15 cm/s group, indicative of increasing temporal asymmetry at 15-30 cm/s compared to slower speeds. The mean iTL for the 30-45 cm/s group was not significantly different from the 15-30 cm/s group. The mean iTL for the > 45 cm/s group was significantly lower than the 30-45 cm/s group, indicating once again that temporal asymmetry of theta cycles initially increases with speed and then plateaus or even decreases slightly at higher speeds (***: p < 0.001, **: p < 0.01,*: p < 0.05).

A similar ANOVA revealed a significant effect of speed on iSkew and iTL as well (iSkew: (F(3,141) = 24, p<0.0001, iTL: F(3,141) = 19.8, p<0.0001). Post hoc Tukey HSD test indicated that the mean iSkew for the 15-30 cm/s group (M = −0.082, SD = 0.11) was significantly more negative than that for the 0-15 cm/s group (M = −0.034, SD = 0.05). However, the mean iSkew for the 30-45 cm/s group (M = −0.088, SD = 0.11) was not significantly different from either the 15-30 cm/s or the >45 cm/s group (M = −0.086, SD = 0.11), confirming that iSkew initially becomes more negative with speed and then saturates at higher speeds. The same post hoc test for iTL showed that mean iTL for the 15-30 cm/s group (M = 0.55, SD = 0.05) was significantly higher than the 0-15 cm/s group (M = 0.53, SD = 0.03). However, the mean iTL for the 30-45 cm/s group (M = 0.55, SD = 0.06) was not significantly different from the 15-30 cm/s group. The mean iTL for the >45 cm/s group (M = 0.54, SD = 0.05) on the other hand was significantly lower than the 30-45 cm/s group. This suggests that iTL also initially increases with speed, then saturates at higher speeds, and then decreases at very fast speeds. These post-hoc tests further indicated that at faster speeds, instantaneous theta amplitude continued to increase, while instantaneous asymmetry remained unchanged. These results show that theta instantaneous amplitude and instantaneous shape have differential relationships to running speed.

## DISCUSSION

The central finding of this work is that theta oscillations in the postrhinal cortex are temporally asymmetric (Figures 2, 3). We have developed an additional method of quantifying temporal asymmetry (instantaneous skewness) that takes the weighted center of mass of each oscillatory cycle into account, resulting in more precise descriptions of the true *shape* of a theta cycle (iSkew; Figure 1). Using this method, we have shown that both the instantaneous amplitude and shape of theta oscillations fluctuate rapidly and mostly independently (Figures 4, 5). This disconnect between the amplitude and shape of theta oscillations is also evident as a function of running speed: while both amplitude and temporal asymmetry of cycles increase with initial increases in speed, the asymmetry of theta cycles plateaus above 25 cm/s while the amplitude continues to increase (Figure 6, 7). These results suggest that there are likely to be partially independent mechanisms controlling the amplitude versus shape of theta cycles in the POR.

We have also shown that postrhinal electrode locations with the highest theta power have, on average, negatively skewed peak-to-peak cycle shapes in the time domain (Figure 3). This means that the majority of cycles at these locations have longer peak-to-trough falling durations, with shorter subsequent trough-to-peak rise times. An important focus of future work will be to repeat these recordings and analyses using laminar probes that sample every layer of the POR at the same time. Such recordings can help to construct current source density (CSD) plots that identify current sinks and sources. The asymmetric falling and rising phases of POR theta cycles can then be aligned to these sinks and sources, estimating the direction of precise current flow associated with the extended falling phase and rapid rising phase of POR theta reported here. Active sinks typically correspond to net synaptic excitation of neurons surrounding the electrode, while active sources correspond to net synaptic inhibition (Buzsaki et al, 2012; Einevoll et al., 2012; Einevoll et al, 2013; Kajikawa & Schroeder, 2011; Ahmed & Cash, 2013). Mechanistically, temporally synchronized inhibition (active sources) onto pyramidal neurons in the vicinity of the electrode during the rapid rising phase may be a critical determinant of the instantaneous temporal asymmetry of the theta cycle: the more synchronized the local inhibition, the faster the rising phase seen in the LFP and the higher the instantaneous temporal asymmetry of that theta cycle. Such precisely synchronized firing of local inhibitory neurons would then be expected to give rise to more precise windows for the synchronization of local excitatory neurons (Pastoll et al., 2013; Hernández-Pérez et al., 2020).

We wish to stress the precise terminology used in this study when discussing temporally asymmetric theta oscillations in the context of previous work on asymmetric waves across disciplines. In statistical analysis, skewness is a measure of the asymmetry of any distribution. We utilize this statistical calculation and definition in this manuscript, with iSkew being a metric that calculates the temporal skewness of each peak-to-peak inverted theta cycle, hence giving a quantitative metric of asymmetry. With these definitions, skewness and asymmetry both refer to the temporal waveform shape, and stating that a theta cycle is temporally skewed means that it has an asymmetric waveform that starts to approach a sawtooth-like shape. In contrast, Bullock et al. (1997), applying the techniques introduced in the context of bispectral analyses of ocean waves by Hasselman et al. (1963), used the terms skewness and asymmetry to refer to two distinct properties of waves. Skewness was defined by them as the “non-equivalance of EEG waves around the horizontal time axis”, while asymmetry was defined as the non-equivalence of “EEG deflections around the vertical or voltage axis”. Our time-domain analysis using iSkew is thus measuring only what Bullock et al. (1997) called “asymmetry”, and not what they called “skewness”. To remove any future ambiguity across LFP waveform shape analyses with roots in statistical distributions versus oceanography, we recommend that the terms “temporal skewness” and “temporal asymmetry” both be used interchangeably to refer to the shape of the oscillation over time (across the vertical axis). Under this definition (as used in this manuscript), a theta cycle that has a sawtooth-like shape is temporally skewed and shows strong temporal asymmetry. We recommend the terms “voltage skewness” and “voltage asymmetry” both be used interchangeably to refer to the shape of the oscillation across the horizontal axis. The term “non-linearity” of EEG or LFP refers to asymmetry around either the horizontal and/or vertical axes (Bullock et al., 1997; Sheremet et al, 2016). Indeed, the non-linearity of theta oscillations (across both horizontal and vertical axes) in the hippocampus has been reported to increase as a function of running speed (Sheremet et al., 2016). While we do not quantify the skewness around the horizontal axis (voltage asymmetry) in this study, our demonstration of temporal asymmetry across the vertical axis is consistent with what was observed by Sheremet et al. (2016) using non-linear measures. To summarize the findings across manuscripts using the unified terminology proposed here, Sheremet et al. (2016) showed that hippocampal theta oscillations, as measured using bispectral analysis, show more temporal asymmetry and more voltage asymmetry as a function of increasing speed. We show, using single cycle analysis, that postrhinal theta cycles become more temporally asymmetric with initially increasing speed but reach a plateau at medium speeds. Future work will need to examine the voltage asymmetry of these postrhinal theta cycles and compare the precise equivalence between single-cycle and bispectral analyses of temporal and voltage asymmetry (Sheremet et al., 2016; Sheremet et al., 2019). This future work will also be needed to quantify how gamma power, gamma frequency, and theta-gamma coupling change as a function of speed and as a function of increasing temporal or voltage asymmetry (see Zhou et al. (2019) for a thorough review and analysis of the methodology necessary to answer these questions in the context of asymmetric theta oscillations in the hippocampus).

Single cycle temporal asymmetry of hippocampal and entorhinal theta cycles has also previously been computed using the asymmetry index and minor variants (Belluscio et al., 2012, Hernández-Pérez et al., 2020). Instantaneous Trough Location (iTL, as used in this manuscript) and the asymmetry index are very similar metrics, as both consider ratios of the ascending and descending parts of the waveform. However, the two metrics differ slightly in the method of extracting individual cycles. Belluscio et. al. (2012) filtered the LFP in the 1-80 Hz band and found the local minima and maxima to estimate cycle bounds. Our observations suggest that this method can be sensitive to noise unless theta power is high. Instead, we use a two-step process to define the bounds of each cycle. First, we find the approximate cycle bounds by filtering the LFP in the 6-10 Hz range and identify the minima and maxima for each cycle. We then refine the precise peaks and troughs closest to these using the 6-40 Hz filtered signal. This allows us to remove ambiguity about the theta cycle bounds while retaining information about the waveform shape. Our second method – instantaneous skewness (iSkew) – is rather different from the asymmetry index (see Figure 1 and Methods) in that it takes the full shape of each cycle into account. It does so by applying the familiar statistical concept of skewness of distributions to individual theta cycles (see above). This iSkew method is able to then estimate the center of mass of each cycle, factoring in precise differences in shape. Although results are qualitatively similar with both iTL and iSkew, iSkew more faithfully captures the true shape of the cycle, and we recommend iSkew as the method of choice for single cycle temporal asymmetry calculations. iSkew can also easily be adapted to calculate single cycle voltage asymmetry in the future.

Unlike hippocampal theta (Buzsaki, 2002), a theoretical framework for the role of theta oscillations in postrhinal spatial computations is currently lacking. Indeed, there is even conflicting evidence regarding the theta-modulation of postrhinal cells. Furtak et al. (2012) recorded LFP together with single-unit activity to show that 38% of postrhinal cells were significantly phase-locked to the theta (predominantly to the trough, whereas putative fast-spiking inhibitory neurons seemed to prefer the peak of theta). LaChance et al. (2019), using only spike trains and not LFP, reported only 1% of their postrhinal cells as showing theta-frequency rhythmicity. While LaChance et al. (2019) mentioned the more caudomedial anatomical location of their postrhinal cells as a potential explanation for this discrepancy, it should be noted that calculating theta rhythmicity from spike-trains is not the same as calculating phase-locking to the theta rhythm seen in the LFP. The relatively low peak firing rate of postrhinal allocentric, egocentric, and conjunctive cells (~7 Hz, LaChance et al., 2019) makes spike-based theta rhythmicity a noisier calculation than phase-locking to the LFP. Thus an important next step towards understanding the role of theta in supporting the encoding of allocentric and egocentric space shown by POR cells (Furtak et al., 2012; LaChance et al., 2019) would be to run the same theta phase-locking analysis of cells recorded at distinct postrhinal anatomical locations to confirm whether they show distinct degrees of phase-locking to theta and whether these differences also correspond to altered temporal waveforms of theta at these distinct locations. In the hippocampus, dorsal CA1 locations show greater temporal and voltage theta asymmetry than intermediate CA1 locations (Sheremet et al., 2016). In the medial entorhinal cortex, temporal asymmetry also decreases across the dorsoventral axis (Hernández-Pérez et al., 2020). Similar anatomical variations may be possible across the postrhinal cortex (Furtak et al., 2007; Agster & Burwell, 2009), potentially with important computational implications for the role of postrhinal theta oscillations in spatial encoding (Maurer et al., 2005; Pastoll et al., 2013; Sheremet et al., 2016; Hernández-Pérez et al., 2020; Lubenov & Siapas, 2009).

## ACKNOWLEDGEMENTS

This work was supported by lab startup funds from the University of Michigan to OJA; the Catalyst Grant from the Michigan Institute for Computational Discovery & Engineering to OJA; NIMH R01MH108729 and NSF IOB-0522220 grants to RDB; and an NIH F32-MH084443 award to SCF. We are also grateful for support from the Center for Consciousness Science at the University of Michigan.

